# Central auditory decline precedes cochlear deficits in a D-galactose mimetic model of aging

**DOI:** 10.64898/2026.05.30.729003

**Authors:** Aravindakshan Parthasarathy, Edward L. Bartlett

## Abstract

Age-related hearing loss reflects a mixture of concurrent peripheral cochlear and central auditory pathway degeneration. Disentangling their relative contributions has remained challenging because both decline together with natural aging. Here, we used systemic D-galactose (D-gal) administration to selectively accelerate central auditory aging while preserving peripheral cochlear function. Eight male Fischer 344 rats received daily subcutaneous D-gal injections (500 mg/kg) for eight consecutive weeks. Two-channel auditory evoked potentials, including auditory brainstem responses (ABRs) and envelope-following responses (EFRs), were recorded at three time points: pre-injection, immediately following injections (2 months post), and 12 months post-injection cessation. Two months post-injection, there was a significant reduction in ABR wave amplitudes from central auditory generators (Waves IV and V) and EFRs to slow modulation frequencies (≤45 Hz), which reflect midbrain and cortical generators. In contrast, ABR thresholds, wave amplitudes from peripheral generators (Waves I and III), and EFRs to fast modulation rates (>128 Hz) reflecting peripheral neural activity remained unchanged at this time point, indicating preserved cochlear and early brainstem function. Peripheral deficits emerged only by the 12-month follow-up, with elevated thresholds, reduced peripheral wave amplitudes, and reduced EFRs at fast modulation rates. These findings demonstrate a clear temporal dissociation in which central auditory decline precedes peripheral cochlear and neural loss in the D-gal model. This paradigm provides a tractable experimental window for studying mechanisms of central auditory aging in isolation from cochlear pathology, bridging an important gap between peripheral and central models of presbycusis.

**Highlights:**

- D-galactose accelerates central auditory aging in Fischer 344 rats.
- Central ABR waves and slow EFRs decline within two months of D-gal exposure.
- Peripheral cochlear deficits emerge only after a 12-month delay.
- Central auditory decline precedes cochlear loss in this model.
- Provides a tractable model to study central auditory aging in isolation.

## Introduction

Age-related hearing loss (ARHL), or presbycusis, is a multifactorial disorder that arises from a complex interplay between peripheral cochlear dysfunction and central auditory pathway degeneration (CHABA, 1988; Helfer et al., 2020; Lentz et al., 2022; Peelle and Wingfield, 2016). At the cochlea, aging leads to outer hair cell stereocilia damage and decreased prestin expression, auditory nerve synapse loss and reductions in endocochlear potential (Cheatham et al., 2004; Ohlemiller et al., 2006; Schmiedt et al., 1996; Sergeyenko et al., 2013). In addition, aging compromises the immune and vascular milieu of the cochlea, altering perilymph composition and cellular homeostasis (Noble et al., 2019; Schmiedt et al., 1996). In parallel, the central auditory system undergoes progressive changes with age, including alterations in excitatory/inhibitory balance, neuronal hyperexcitability, reduced synaptic density, and loss of neuronal volume within auditory subcortical and cortical regions (Bartlett et al., 2024; Caspary et al., 2008; Harris et al., 2022; Herrmann et al., 2016; Herrmann and Butler, 2021; Koehler et al., 2023; Rabang et al., 2012; Robinson et al., 2019) Together, these changes shape the perceptual manifestations of ARHL, such as difficulty understanding speech in noise.

While peripheral damage with age can be selectively modeled using noise exposure or ototoxic drugs (Bielefeld, 2013, 2012; Fernandez et al., 2015; Kujawa and Liberman, 2009), recapitulating central auditory aging in isolation has remained a challenge. Central changes are often secondary to peripheral deafferentation, making it difficult to dissociate cause from consequence. In young animals, both the cochlea and brain are healthy, whereas in aged animals, both are concurrently, but often independently degenerated (Märcher-Rørsted et al., 2026). This covariance makes it difficult to separate central mechanisms of auditory aging from those driven by peripheral deafferentation. Thus, identifying models that selectively accelerate central degenerative processes without concurrent cochlear injury is critical for understanding how aging alone alters auditory neural coding in the central auditory pathway.

The aging brain itself undergoes widespread cellular stress. Oxidative damage, mitochondrial dysfunction, and declines in ATP production alter neuronal and glial viability and promote microglial activation and neuroinflammatory signaling (Helfer and Jesse, 2021; Watson et al., 2017; White and Someya, 2023). One established approach to accelerate these processes is via systemic administration of D-galactose (D-gal), a reducing sugar that, at supraphysiological doses, induces oxidative stress, mitochondrial DNA damage, and advanced glycation end-product accumulation (Chen et al., 2010; Ho et al., 2003). Chronic D-gal exposure produces phenotypes that mimic natural aging in multiple tissues, including the cerebral cortex and cochlea (Chen et al., 2010; Guo et al., 2020; Peng et al., 2023). However, the temporal sequence of auditory effects following D-gal treatment remains underexplored.

Here, we establish systemic D-galactose administration as a mimetic model of central auditory aging. Using non-invasive auditory evoked potentials, we demonstrate that systemic D-gal produces accelerated aging phenotypes in central auditory nuclei while preserving peripheral cochlear function during the early phase of exposure. Peripheral cochlear loss emerges only after a time delay, providing a unique temporal window in which the central auditory effects of aging can be studied in isolation from peripheral degeneration. This paradigm offers a tractable approach to dissect the mechanisms of central auditory aging and their contribution to suprathreshold hearing deficits.

## Methods

### Subjects

Eight male Fischer 344 (F344) rats (3-6 months old) were used in this study. Animals were housed in standard laboratory conditions under a 12 h:12 h light-dark cycle with ad libitum access to food and water in a low-noise environment. All procedures were approved by the Purdue Animal Care and Use Committee (PACUC #1111000280) and followed NIH guidelines for the care and use of laboratory animals.

### D-galactose administration

Rats received daily subcutaneous injections of D-galactose (500 mg/kg) dissolved in 1 mL of sterile saline for eight consecutive weeks. This dosing regimen was based on previous aging model studies (Chen et al., 2010) and is known to induce systemic oxidative stress and central neurodegeneration.

### Auditory Evoked Potential Recordings

Two-channel auditory evoked potential (AEP) recordings were obtained at three time points: pre-injection (baseline), immediately after completion of injections (2 months post-injection), and 12 months post-injection cessation. Procedures followed established, published protocols (Lai et al., 2017; Parthasarathy et al., 2014; Parthasarathy and Bartlett, 2012). Briefly, animals were anesthetized with 1.8-2% isoflurane and subdermal needle electrodes (Ambu) were inserted. Channel 1 (Fz to Cz, mid-sagittal) emphasized peripheral generators; Channel 2 (C3 to C4, interaural line) emphasized central generators. The reference electrode was placed under the mastoid of the right ear (ipsilateral to the speaker) and a ground electrode on the back of the animal. Electrode impedances were below 1 kΩ (RA4LI, TDT). Animals were then sedated with intramuscular dexmedetomidine (0.2-0.3 mg/kg, Dexdomitor), and recordings were performed 10-15 min after isoflurane removal to avoid anesthetic effects, allowing approximately 2-3 hours of recording time. Stimuli were presented free-field to the right ear (90° azimuth) using a calibrated Bowers & Wilkins speaker positioned 115 cm away.

### Auditory Brainstem Responses (ABRs)

ABRs were elicited by clicks (0.1 ms) and tone pips (1-32 kHz, 2 ms total duration with 0.5 ms cos^2^ rise-fall) presented in alternating polarity at 26.6 Hz (1500 repetitions). Stimuli ranged from 95 to 15 dB pSPL in 10 dB steps. A 20 ms acquisition window was used, with band-pass filtering from 30 Hz to 3 kHz.

ABR thresholds were visually identified as the lowest level eliciting a repeatable waveform and independently confirmed by two additional blinded researchers. As established previously (Parthasarathy and Bartlett, 2012), Wave I and Wave III were quantified from Channel 1, which emphasizes peripheral neural generators on the auditory nerve and brainstem, and Waves IV and V were quantified from Channel 2, which emphasizes central auditory generators in the midbrain.

### Envelope-Following Responses (EFRs)

EFRs were collected in the same session following ABRs using sinusoidally amplitude-modulated (AM) Gaussian noise carriers (200 ms duration, 5ms onset/offset cosine ramp). AM frequencies ranged from 16 to 2048 Hz in ½-octave steps, with 200 repetitions per condition and a 300 ms recording window. FFT analysis in MATLAB extracted response amplitudes at the modulation frequency, as described previously (Lai et al., 2017; Parthasarathy et al., 2010; Parthasarathy and Bartlett, 2011). Channel 1 was most sensitive to high modulation frequencies (90-2048 Hz), whereas Channel 2 captured low modulation frequencies (8-90 Hz) (Lai et al., 2017; Parthasarathy and Bartlett, 2012). A composite temporal modulation transfer function was therefore constructed by combining EFRs from the two channels at their respective sensitive modulation frequencies.

### Statistical Analysis

All analyses were performed in MATLAB. Because all animals received D-galactose injections, the study employed a within-subject longitudinal design; the pre-injection recording served as the baseline comparator. One animal lacked data at the 12-month time point and was excluded from three-timepoint analyses; additional animals were excluded for individual measures with missing data. Sample sizes for each analysis are provided in Table 1. Greenhouse-Geisser correction was used for sphericity violations.

**Table 1.** Summary of two-way repeated-measures ANOVA results for ABR thresholds, ABR wave amplitudes (Waves I, III, IV, and V), and EFR amplitudes. All p-values are Greenhouse-Geisser (GG) corrected for sphericity. ABR wave analyses were conducted over sound levels 55-95 dB SPL; EFR analyses were conducted over modulation frequencies 16-2048 Hz. Significant post-hoc pairwise comparisons (Tukey’s HSD) are indicated by asterisks in Figures 1 and 2.

| Measure | n | Factor | df | F | p |
| --- | --- | --- | --- | --- | --- |
| ABR Thresholds | 7 | Time | 2, 12 | 31.96 | <0.001 |
|  |  | Frequency | 6, 36 | 650.52 | <0.001 |
|  |  | *Time × Frequency* | 12, 72 | 8.26 | 0.002 |
| ABR Wave I | 7 | Time | 2, 12 | 20.64 | <0.001 |
|  |  | Sound level | 1, 6 | 324.34 | <0.001 |
|  |  | *Time × Sound level* | 2, 12 | 17.98 | 0.001 |
| ABR Wave III | 7 | Time | 2, 12 | 25.47 | <0.001 |
|  |  | Sound level | 1, 6 | 746.53 | <0.001 |
|  |  | *Time × Sound level* | 2, 12 | 22.77 | <0.001 |
| ABR Wave IV | 6 | Time | 2, 10 | 25.68 | <0.001 |
|  |  | Sound level | 1, 5 | 199.58 | <0.001 |
|  |  | *Time × Sound level* | 2, 10 | 24.35 | <0.001 |
| ABR Wave V | 6 | Time | 2, 10 | 19.25 | 0.006 |
|  |  | Sound level | 1, 5 | 110.77 | <0.001 |
|  |  | *Time × Sound level* | 2, 10 | 18.64 | 0.006 |
| EFR | 5 | Time | 2, 8 | 31.77 | 0.001 |
|  |  | Modulation frequency | 1, 4 | 393.57 | <0.001 |
|  |  | *Time × Modulation frequency* | 2, 8 | 13.44 | 0.004 |

Two-way repeated-measures ANOVAs tested within-subject effects of Time (Pre, 2 months, 12 months) and a second factor (Tone frequency for ABR thresholds; Sound Level for ABR wave amplitudes; Modulation frequency for EFRs), along with their interaction. All pairwise post-hoc comparisons used Tukey’s HSD correction. Statistical significance was set at α = 0.05. Main effects are summarized in Table 1; significant post-hoc comparisons are indicated by asterisks in the corresponding figures.

## Results

### D-galactose selectively impairs central, but not peripheral, auditory function

To examine whether systemic D-galactose (D-gal) administration differentially affects peripheral versus central auditory processing, ABRs were recorded at three time points: prior to D-gal exposure (Pre), immediately after eight weeks of daily injections (2 months post), and 12 months post-injection cessation (Fig. 1A). Given their age preceding D-gal exposure, animals at the final time point would be expected to express a middle-aged phenotype with increased ABR thresholds (Parthasarathy et al., 2014), and minimal changes in IC responses or GABAergic markers (Caspary et al., 2008; Syka, 2010).

**Figure 1.**
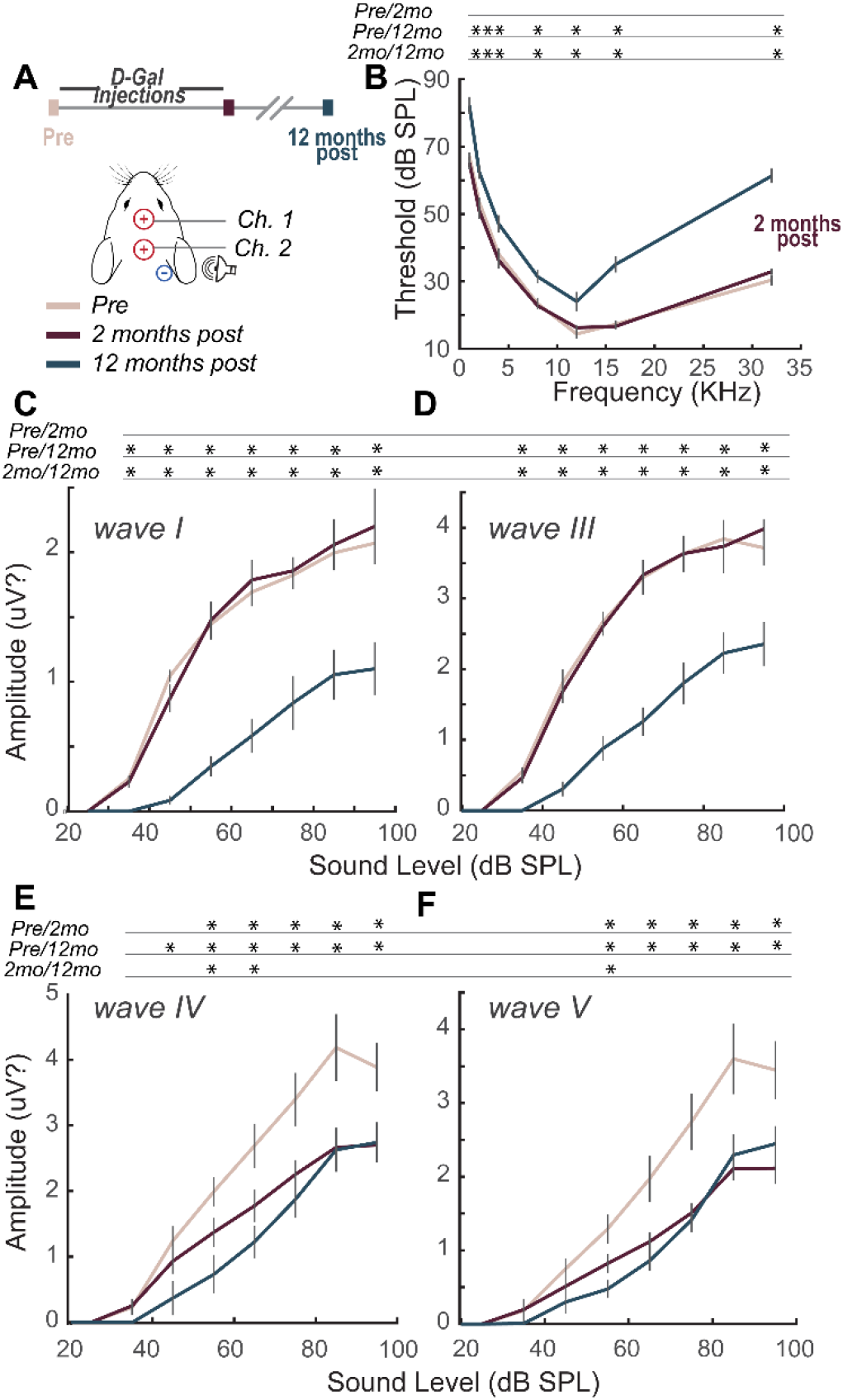
D-galactose induces early central auditory dysfunction with delayed peripheral loss. (A) Timeline of D-gal injections and recording sessions. (B) Mean ABR thresholds across frequencies at baseline (Pre), 2 months post-injection, and 12 months post-injection. (C–F) ABR amplitude-level functions for Waves I, III, IV, and V. Asterisks indicate significant post-hoc pairwise differences (Tukey’s HSD, p < 0.05). Early reductions in Waves IV and V amplitudes indicate central auditory deficits preceding peripheral deterioration. Error bars represent ± SEM.

ABR thresholds showed a significant main effect of Time and a significant Time × Tone frequency interaction (Table 1). Post-hoc comparisons revealed that thresholds at 12 months post were significantly elevated relative to both the Pre baseline (mean difference = 13.9 dB, p = 0.003) and the 2 months time point (mean difference = 14.9 dB, p = 0.001), whereas thresholds did not differ significantly between Pre and 2 months post (mean difference = 1.0 dB, p = 0.80, Fig. 1B). These results indicate no acute cochlear dysfunction immediately following the D-gal injection period, with threshold elevation emerging only at the 12-month follow-up.

ABR wave amplitude analyses also revealed significant main effects of Time, Sound Level, and their interaction for all four waves (Table 1). For peripheral generators, Wave I and Wave III showed no significant amplitude difference between Pre and 2 months post, but both exhibited significant reductions at 12 months relative to Pre and 2 months post (Fig. 1C, D), consistent with delayed peripheral deafferentation.

In contrast, Waves IV and V, generated by more central auditory nuclei, showed a critically different temporal pattern (Fig. 1E, F). Significant amplitude reductions were observed between Pre and 2 months post at multiple sound levels, with further declines at 12 months post relative to Pre across all tested levels. Wave V (Fig. 1F) showed the same pattern, with significant reductions at 2 months relative to Pre, and further reductions at 12 months.

Together, these results demonstrate a clear temporal dissociation, with central ABR wave amplitudes (Waves IV and V) declining early, before any threshold or peripheral wave changes, indicating selective vulnerability of central auditory circuits to D-gal–induced oxidative stress. Peripheral deficits emerge only later, after prolonged post-treatment aging.

### Temporal processing deficits measured by EFRs follow similar patterns of central vulnerability

EFR amplitudes were assessed across a wide range of amplitude modulation frequencies, with stimuli presented at 30 dB above each animal’s individual ABR threshold to equalize sensation level. Despite this normalization, EFR amplitudes exhibited the same temporal pattern of decline observed in ABRs. A two-way repeated-measures ANOVA revealed significant main effects of Time and Modulation Frequency, and a significant Time × Modulation Frequency interaction (Table 1), indicating that the time course of EFR change differed across the modulation rate spectrum.

Specifically, EFRs to rapid modulation rates (>300 Hz), which are dominated by peripheral neural generators such as the auditory nerve and cochlear nucleus, remained stable immediately after the eight-week D-gal injection period but showed significant reductions at the 12-month follow-up (Fig. 2), similar to older animals (Parthasarathy and Bartlett, 2012). In contrast, EFRs to slow modulation rates (<128 Hz), which reflect central auditory cortical generators, were significantly reduced immediately after eight weeks of D-gal administration and remained depressed one year later (Fig. 2).

**Figure 2.**
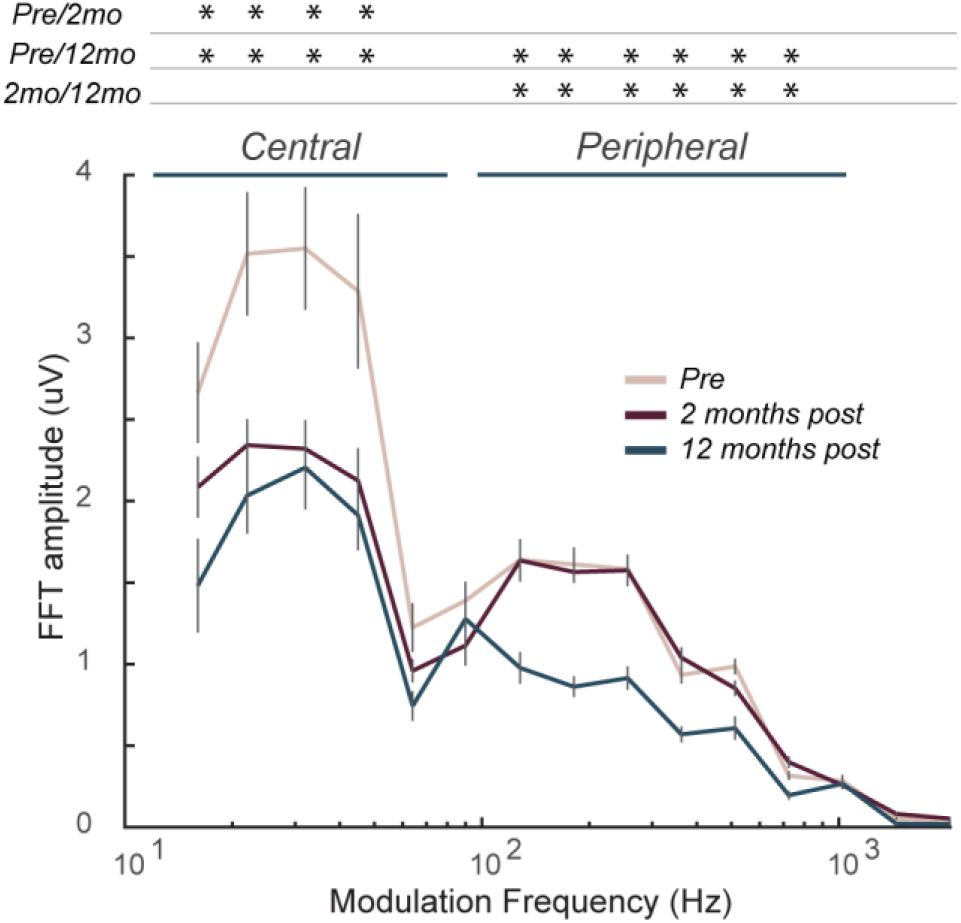
Temporal processing deficits following D-galactose administration mirror the time course of central auditory decline. Mean envelope-following response (EFR) amplitudes (µV) across modulation frequencies measured before (Pre), 2 months after the start of daily D-galactose injections, and 12 months post-injection. Stimuli were presented at 30 dB above individual ABR thresholds to equalize sensation levels. Asterisks indicate significant post-hoc pairwise differences (p < 0.05). EFRs to low modulation rates (≤45 Hz) were significantly reduced at both 2 months and 12 months relative to Pre. EFRs to higher modulation rates (> 128 Hz) were unaffected at 2 months but declined significantly by 12 months. Error bars represent ± SEM.

These findings confirm that D-gal-induced oxidative stress preferentially and rapidly disrupts central auditory temporal coding, even when peripheral neural synchrony and cochlear thresholds are preserved. Over longer time scales, peripheral contributions also deteriorate, mirroring the delayed ABR amplitude loss (Fig. 1).

## Discussion

Aging of the auditory system is a multifaceted process involving both peripheral cochlear degeneration and deterioration of the central auditory pathway (Helfer et al., 2020; Helfer and Jesse, 2021). While peripheral damage can be modeled effectively using noise exposure or ototoxic agents (Ohlemiller et al., 2000; Schmiedt et al., 2002), experimental paradigms that isolate central auditory aging in the absence of cochlear injury have remained elusive.

In this study, we demonstrate that systemic D-galactose (D-gal) administration provides a tractable model to selectively accelerate central auditory aging. Following eight weeks of D-gal exposure, animals exhibited a biphasic pattern of auditory decline. The initial phase was characterized by central auditory dysfunction, reflected in reduced ABR Waves IV and V amplitudes and diminished EFR synchrony at modulation frequencies ≤45 Hz AM, both indices of degraded neural coding within midbrain and cortical circuits. Importantly, these deficits occurred without concomitant changes in hearing thresholds, peripheral ABR waves (I and III), or EFRs for modulation rates above 300 Hz, indicating preserved cochlear and early brainstem function. Peripheral declines emerged over the subsequent 12 months, indicated by elevated thresholds, reduced Wave I and III amplitudes, and loss of EFR amplitudes for modulation rates above 300 Hz, suggesting that central deterioration precedes functional peripheral aging in the D-gal model.

The EFR paradigm was particularly valuable for dissociating central and peripheral contributions because its temporal modulation rate selectively emphasizes distinct neural generators. Prior work has shown that EFRs at high modulation rates (∼1000 Hz) are sensitive to auditory nerve dysfunction, whereas EFRs at slow modulation rates (∼40 Hz AM) reflect cortical activity and are reduced by anesthesia (McHaney et al., 2024; Parthasarathy and Bartlett, 2012; Parthasarathy and Kujawa, 2018; Shaheen et al., 2015; Zink et al., 2024). The current findings extend this framework by demonstrating that D-gal-induced oxidative stress selectively disrupts slow modulation rate EFRs early, consistent with central vulnerability to metabolic compromise.

Future studies will aim to resolve the precise temporal emergence of peripheral deficits between 2 months and 12 months and to perform histological validation in both cochlear and central auditory structures. Such analyses will help define the mechanisms underlying D-gal-induced degeneration, and dissociate mitochondrial, glial, and/or vascular dysfunction, and establish the specific relationship between functional changes and cellular loss or synaptic dysfunction.

In conclusion, systemic D-galactose administration produces a temporally specific model of auditory aging in which central neural decline precedes peripheral loss. This model provides a valuable experimental window to study the mechanisms of central auditory aging and maladaptive plasticity independently of cochlear pathology, thereby bridging a critical gap between peripheral and central models of presbycusis.

## Acknowledgements

The authors thank Julie Ann Luna Torres and Charneka Hopkins for assistance with data collection and Audrey Transue for assistance with statistical analysis. This work was funded by the Department of Defense (W81XWH2110602 to ELB and AP).

## CRediT author contributions

AP: Conceptualization, Methodology, Investigation, Formal analysis, Writing - original draft, Writing - review & editing, Funding acquisition. ELB: Conceptualization, Methodology, Resources, Supervision, Writing - review & editing, Funding acquisition.

## Data availability

Data supporting the findings of this study are available from the corresponding author upon reasonable request.

